# Semisynthetic pH-sensitive fluorophores for imaging exocytosis and endocytosis

**DOI:** 10.1101/118505

**Authors:** Magalie Martineau, Agila Somasundaram, Jonathan B. Grimm, Todd D. Gruber, Daniel Choquet, Justin W. Taraska, Luke D. Lavis, David Perrais

## Abstract

The GFP-based superecliptic pHluorin (SEP) enables detection of exocytosis and endocytosis, but its performance has not been duplicated in red fluorescent protein scaffolds. Here we describe ‘semisynthetic’ pH-sensitive protein conjugates that match the properties of SEP. Conjugation to genetically encoded self-labeling tags or antibodies allows visualization of both exocytosis and endocytosis, constituting new bright sensors for these key steps of synaptic transmission.

---

Synaptic transmission is mediated by the rapid fusion of synaptic vesicles (SVs) with the plasma membrane. Precise monitoring of exocytosis is important for elucidating fundamental mechanisms of cell–cell communication, and investigating the underlying causes of neurological disorders^1^. The lumen of synaptic vesicles are typically acidified (pH 5.6) by the action of vesicle-resident V-ATPases, which creates the driving force for neurotransmitter uptake. Upon fusion with the plasma membrane, the contents of the vesicle rapidly equilibrate with the extracellular environment (pH 7.4). This large change in pH allows for the visualization of exocytosis using a pH-sensitive variant of green fluorescent protein (GFP) that is expressed as a fusion with a vesicular membrane protein^2^. This ‘superecliptic pHluorin’ (SEP) exhibits ideal properties for detecting the change in pH upon vesicle fusion, with near-ideal pKa, cooperative protonation, and low background fluorescence in the protonated state, making it an excellent tool for monitoring exocytosis in living cells^2^.

A useful extension of this technology has been the creation of pH sensors based on red fluorescent proteins (RFPs) such as mOrange2^3^, pHTomato^4^, pHoran4^5^, and pHuji^5^. Longer excitation wavelengths are less phototoxic, elicit lower levels of autofluorescence, facilitate multicolor imaging experiments, and allow concomitant use of optogenetics. Nevertheless, it has proven difficult to engineer red-shifted pHluorins that match the optimal p*K*a, cooperativity, and dynamic range of SEP, perhaps due to inherent limitations in RFP scaffolds. More generally, these techniques rely on overexpression of reporter proteins in SVs and the effect of overexpression is a confounding factor in interpreting experimental results^6^.

Given the limitations of pH-sensitive RFPs and the potential problems with overexpression of sensor proteins, we pursued an alternative strategy: creation of ‘semisynthetic’ pH indicators using organic pH-sensitive dyes attached to either expressed self-labeling tags such as the SNAP-tag^7^ or antibodies that recognize native vesicular proteins (**Fig. 1a**). To match the performance of SEP, we required a pH-sensitive organic dye that can undergo a cooperative transition from a bright, fluorescent form at neutral pH to a nonfluorescent form at low pH. Unfortunately, the majority of pH-sensitive dyes do not meet these requirements. The archetypical small molecule pH-sensor is fluorescein (Fl, **1**, **Fig. 1b**), which transitions between a highly fluorescent dianion (**1^2–^**) and a less fluorescent monoanion (**1^−^**) with a relatively low pKa value of 6.3 (**Fig. 1c**)^8^. Other unsuitable synthetic pH probes include the ratiometric seminapthorhodofluor (SNARF) dyes^9^ that exhibit high background, as well as cyanine and rhodamine-based pH sensors that show the opposite pH sensitivity profile^10,11^.

**Figure 1.**
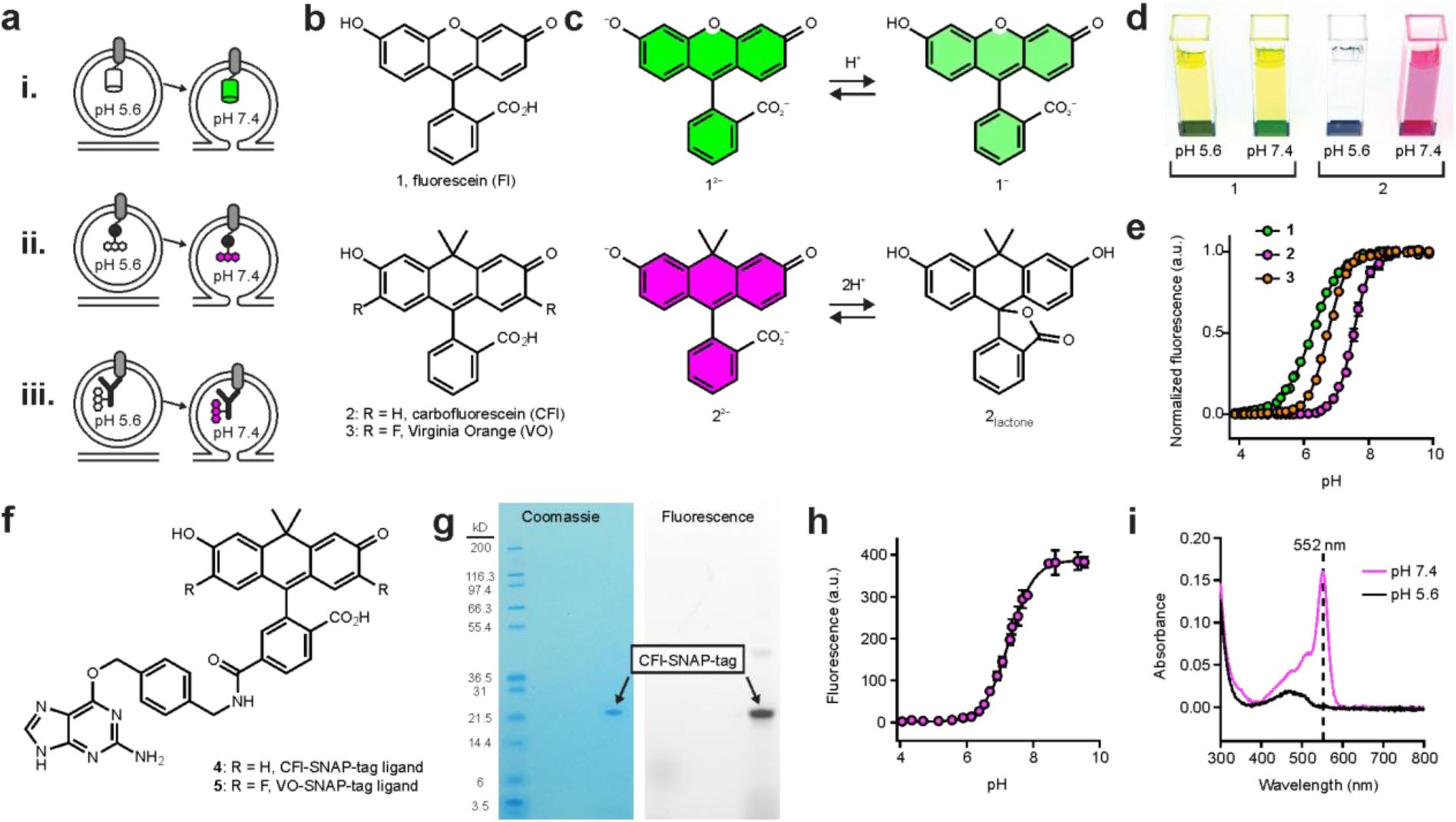
Design and characterization of semisynthetic fluorescent reporters for exocytosis. **(a)** Three protein labeling strategies for imaging exocytosis. Cartoons depicting vesicles expressing pH-sensitive fluorescent protein, such as SEP or pHuji, fused to a SV protein (i) or SNAP-tag enzyme fused to an SV protein and labeled with a pH-sensitive organic fluorophore (ii) and (iii) labeling of endogenous SV proteins with an antibody conjugated to a pH-sensitive dye. (**b**) Chemical structures of Fl (**1**), CFl (**2**), and VO (**3**). (**c**) Fl and CFl respond differently to pH. Fluorescein undergoes noncooperative protonation from the dianion (**1^2−^**) to the monoanion (**1^−^**) whereas carbofluorescein undergoes cooperative protonation from the highly colored dianion (**2^2−^**) to a colorless lactone form (**2_lactone_**). (**d**) Image of cuvettes containing compounds **1** or **2** (5 μΜ) at pH 5.6 and pH 7.4. (**e**) Plot of fluorescence *versus* pH for compounds **1**, **2** and **3**. (**f**) Chemical structure of CFl–SNAP-tag ligand (**4**) and VO–SNAP-tag ligand (**5**). (**g**) PAGE of SNAP-tag protein labeled with compound **4** and visualized by Coomassie staining or by fluorescence. (**h**) Plot of fluorescence *versus* pH for **4**–SNAP-tag conjugate. Found pKa = 7.3 and ηH = 1.2. (**i**) Absolute absorbance of **4**-SNAP-tag conjugate (˜3 μΜ) at pH 7.4 (magenta) and pH 5.6 (black).

We recently synthesized new derivatives of fluorescein (**1**) where the xanthene oxygen was replaced with a gem-dimethylcarbon moiety. This work resulted in ‘carbofluorescein’ (CFl, **2**, **Fig. 1b**)^12^, and the difluorinated derivative ‘Virginia Orange’ (VO, **3**, **Fig. 1b**)^13^. We discovered that this oxygen→carbon substitution elicited significant changes in photophysical and chemical properties of the fluorescein scaffold. Fl (**1**) exhibits λex/λem = 491 nm/510 nm at high pH, whereas CFl (**2**) and VO (**3**) are red-shifted with λex/λem = 544 nm/567 nm and 555 nm/581 nm, respectively. In addition to this bathochromic shift, the pH sensitivity of the dyes was markedly different. Fluorescein exhibits strong visible absorption at both pH 5.6 (vesicle pH) where the monoanion **1^−^** dominates, and pH 7.4 (extracellular pH) where the dianion form **1^2–^** is prevalent—this can be observed by eye (**Fig. 1c,d**). In contrast, CFl (**2**) undergoes a cooperative transition between a highly colored dianion species (**2^2–^**) and a colorless lactone form (**2_lactone_**; **Fig. 1c**). This is also evident visually as a solution of CFl (**2**) is colorless at pH 5.6, but shows robust visible absorption at pH 7.4 (**Fig. 1d**). Fluorescence-based titrations (**Fig. 1e**) gave p*K_a_* values of 6.3 and Hill coefficient (ηΗ) value of 0.97 for fluorescein (**1**), consistent with previous reports^8^. CFl (**2**) and VO (**3**) displayed p*K_a_* values of 7.5 and 6.7, and ηΗ values of 1.6 and 1.5, respectively. This cooperative transition likely stems from the altered lactone–quinoid equilibrium observed in the carbon-containing analogs of fluorescein and rhodamine dyes^12^. The longer absorption and emission wavelengths, higher p*K_a_*, and the cooperative colorless→colored transition upon increased pH make both CFl and VO attractive scaffolds for building indicators to monitor synaptic vesicle fusion events.

To allow for specific labeling of expressed proteins, we prepared the SNAP-tag ligands attached to CFl (**4**) or VO (**5**) (**Fig. 1f**, **Supplementary Note**). We tested the effects of protein conjugation on the properties of the dye by labeling SNAP-tag protein *in vitro* with CFl–SNAP-tag ligand **4** (**Fig. 1g**). We observed a shift in p*K_a_* to 7.3, and a decreased Hill coefficient (ηΗ = 1.2; **Fig. 1h**). The active site of the SNAP-tag enzyme is flanked with two Lys, one Arg, and one His (PDB structure 3KZZ, DOI: 10.2210/pdb3kzz/pdb). The resulting Coulombic interaction between these positively charged amino acid residues and the CFl label most likely explains the decrease in p*K_a_* upon conjugation^8^. This polar surface might also stabilize the open form of the dye, resulting in the decreased cooperativity of the colored-colorless transition. Despite this lower p*K_a_* value and Hill coefficient, the fluorescence of the SNAP-tag-CFl conjugate is still completely suppressed at pH 5.6 (**Fig. 1i**).

Next, we tested these SNAP-tag-based probes in living cells. Building on existing SEP-based constructs^1,2^ we designed two SNAP-tag fusion proteins: (i) SNAP-tag inserted within an intra-luminal loop of the vesicle acetylcholine transporter VAChT (VAChT-SNAP), and (ii) SNAP-tag protein attached to the luminal C-terminal side of the vesicle protein VAMP2 (VAMP2-SNAP). These proteins were expressed in neuroendocrine PC12 cells, where VAChT is targeted to small synaptic-like vesicles (SSLV) while VAMP2 is found in both SSLV and large dense core vesicles^14,15^. We found that the propensity of the CFl and VO fluorophores to adopt the neutral lactone form (**Fig. 1c,d**) allows for efficient intracellular labeling with SNAP-tag ligands **4** or **5** without the use of other masking groups (*e.g*., acetate esters), which are typically required for fluorescein-based compounds (**Fig. 2a**). To monitor exocytosis, we depolarized cells with stimulation buffer containing high [K^+^] and imaged single small vesicles as they fused with the plasma membrane using total internal reflection fluorescence (TIRF) microscopy. Cells containing VAChT constructs displayed events at high frequency. Events detected in cells co-expressing VAChT-SEP and VAChT-SNAP (labelled with CFl ligand **4**) showed comparable fold increases in fluorescence at exocytosis (2.19 ± 0.07 *vs*. 2.40 ± 0.12) with similar decay kinetics in both the green and red channels (**Fig. 2b,c**). We also compared VAChT-SEP to VAChT-SNAP-VO (**Fig. 2d,e**) and VAChT-pHuji (**Fig. 2f,g**). Like the semisynthetic indicator from CFl ligand **4**, the VAChT-SNAP-VO derived from compound **5** also showed comparable performance to the SEP sensor (2.65 ± 0.10 vs. 2.60 ± 0.16 fold increase; **Fig. 2d,e**). However, in PC12 cells the RFP-based VAChT-pHuji sensor showed lower relative performance when compared with VAChT-SEP under the same conditions (2.01 ± 0.05 vs. 1.32 ± 0.02 fold increase; **Fig. 2f,g**) making events harder to detect with pHuji than with the other pH-sensitive proteins. We also observed individual fusion events using VAMP2-SEP or VAMP2-SNAP-CFl (**Supplementary Fig. 1**), albeit at low frequency, perhaps due to poor incorporation of this construct in PC12 cells.

**Figure 2:**
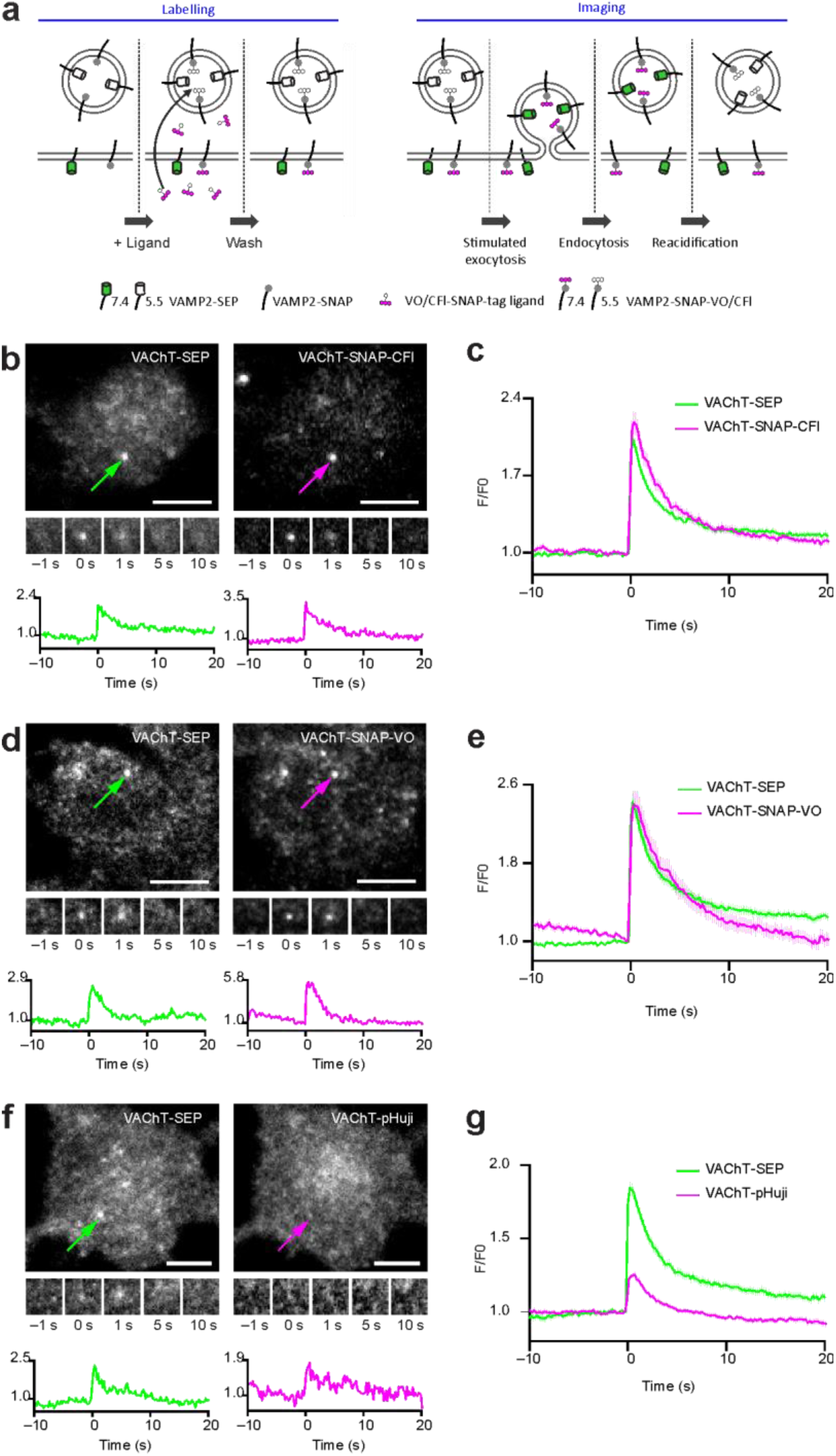
Detection of single exocytosis events in PC12 cells. **(a)** Experimental scheme: prior to imaging, cell-permeable VO/CFl-SNAP-tag ligand is incubated with transfected neuronal cultures to specifically label VAMP2-SNAP. Both VAMP2-SEP and VAMP2-SNAP-VO/CFl are quenched at acidic pH and exhibited maximal fluorescence at the surface. (**b-f**) TIRF microscopic images of PC12 cells expressing VAChT-SEP (left) and VAChT-red shifted indicator (right) undergoing fusion. Images of whole cell with a fusion event indicated with arrow (top); time-lapse of the marked exocytic event (middle, 3.36 × 3.36 pm); normalized intensity traces for the marked event in the green and red channels (bottom); scale bars: 5 μm. (**f**) VAChT-SNAP-CFl. (**g**) VAChT-SNAP-VO. (**h**) VAChT-pHuji. (**g**,**i,k**) Plot of normalized fluorescence versus time averaged across single-vesicle fusion events for vesicles labeled with VAChT-SEP (green) or VAChT-red-shifted indicator (magenta); error bars show ± s.e.m. (**i**) VAChT-SNAP-CFl, n = 113, from 5 cells. (**j**) VAChT-SNAP-VO, n = 116, from 10 cells. (**k**) VAChT-pHuji, n = 126, from 3 cells.

We then tested these sensors in living neurons, focusing on VAMP2 based constructs, which have been used extensively to follow SV exocytosis in neurons^1^. We co-transfected hippocampal neurons with VAMP2-SEP and either VAMP2-pHuji or VAMP2-SNAP incubated with CFl ligand **4** or Virginia Orange ligand **5**. For all the sensors, we observed a robust increase in fluorescence following electrical stimulation in fields covered with transfected axons, signaling SV exocytosis. The relative increase in fluorescence upon SV exocytosis was slightly higher for the SEP channel relative to the red-shifted fluorescent indicators, VAMP2-SNAP-VO (**Fig. 3a, b**), VAMP2-SNAP-CFl (**Supplementary Fig. 2a,b**), and VAMP2-pHuji (**Supplementary Fig. 2c,d**), which behaved similarly. The kinetics of decay, which tracks endocytosis and re-acidification of the vesicle, were similar for all four labels (**Fig. 3b** and **Supplementary Fig. 2b,d**). We also tested if the semisynthetic pH sensor system could be used in multicolor imaging experiments with GFP-based indicators. We co-transfected neurons with the GCaMP6f^16^ and VAMP2-SNAP, which we labeled with CFl ligand **4**. This allowed simultaneous imaging of both calcium ion transients and vesicle fusion in the same cell (**Supplementary Fig. 2e-h**).

**Figure 3.**
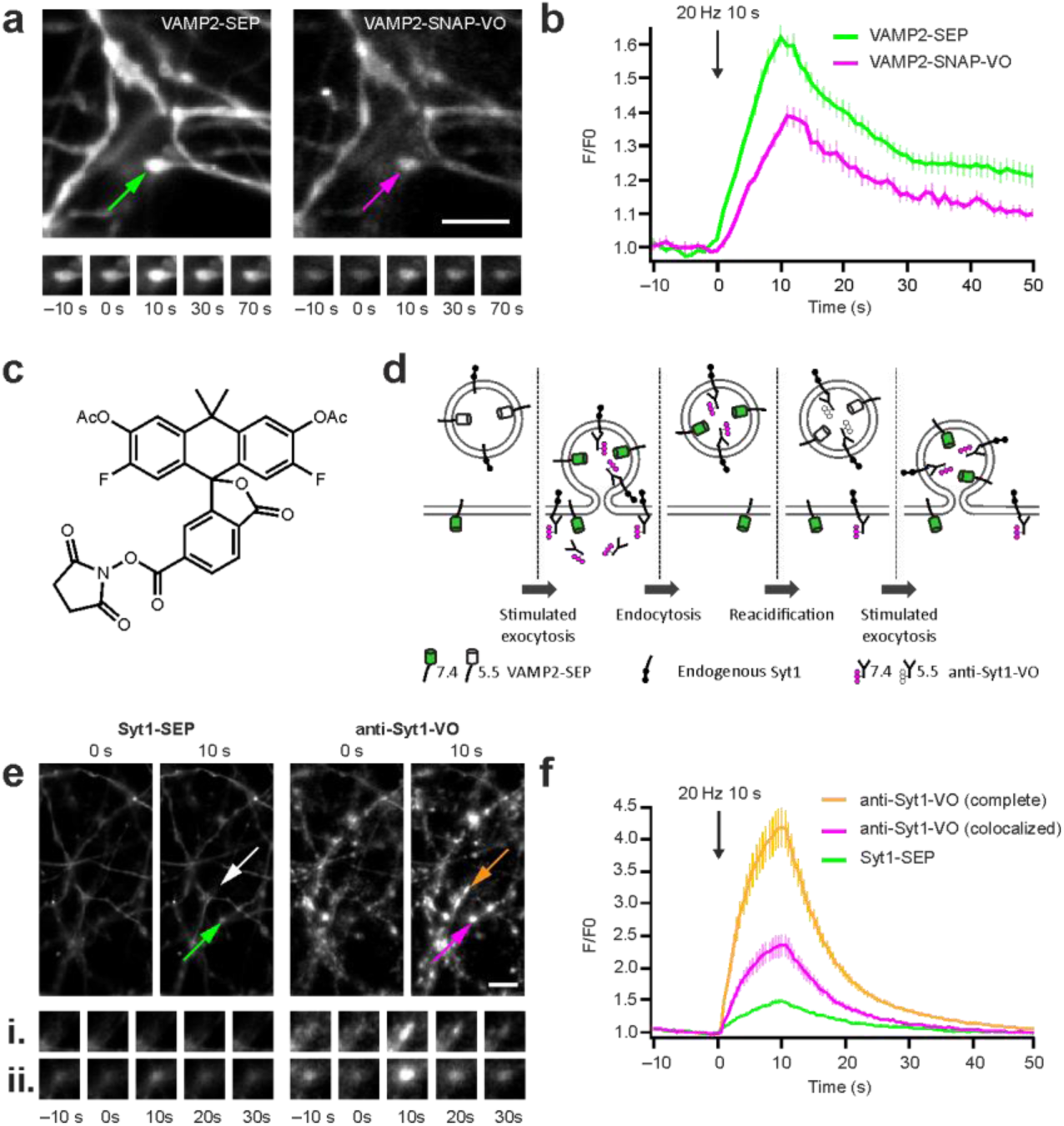
Detection of synaptic vesicle exocytosis and recycling in hippocampal neurons. **(a)** Fluorescence images of hippocampal axons expressing VAMP2-SEP and VAMP2-SNAP-VO with the locations of vesicle fusion indicated with arrows (top); scale bar: 5 μm. Time-lapse images of the exocytosis events (bottom). (**b**) Average VAMP2-SEP (green) and VO–SNAP–VAMP2 (magenta) fluorescence signals in response to field stimulation for 10 s at 20 Hz.; n = 13; data are represented as mean ± s.e.m. Fluorescence decay after stimulation was not different between the two probes (VAMP2-SEP 13.2 ± 1.7 s; VO–SNAP–VAMP2 13.0 ± 1.5 s, *p* = 0.51) (**c**) Chemical structure of VO-NHS ester (**6**). (**d**) Protocol for staining with anti-Syt1-VO. (**e**) Fluorescence images of hippocampal axons expressing Syt1-SEP and stained with anti-Syt1-VO with the locations of vesicle fusion indicated with arrows (top); scale bar: 5 μm. Bottom, time-lapse images of the exocytosis events in boutons containing only anti-Syt1-VO (i) or containing both Syt1-SEP and anti-Syt1-VO (ii). Scale bar: 5 μm. (**f**) Average anti-Syt1-VO (magenta) and Syt1-SEP (green) fluorescence signals colocalized in the same boutons, and average anti-Syt1-VO fluorescence signals present in the complete neuronal population (orange) in response to field stimulation for 10 s at 20 Hz. n = 41. Data are represented as mean ± s.e.m.

Finally, we asked whether SV exocytosis could be detected without overexpression of a reporter protein. To enable imaging of endogenous vesicular proteins, we labeled a monoclonal antibody that recognizes a luminal epitope of synaptotagmin 1 (Syt1), a SV protein, with VOAc_2_-NHS ester (**6**) followed by mild deprotection of the acetate esters using hydroxylamine (**Fig. 3c, Methods**). This antibody has previously been used to detect endogenous Syt1 present on the plasma membrane after exocytosis in active synapses^17^. To mark vesicular Syt1, we incubated neurons with this antibody-VO conjugate (10 nM) for 3 hours in stimulation buffer, followed by extensive washing to remove the extracellularly bound protein conjugate (**Fig. 3d**).

The antibody labelling was done in neurons transfected with Syt1-SEP or VAMP2-SEP to compare the performance of this labeling technique with the overexpressed, genetically encoded GFP-based pH sensors. Electrical stimulation evoked a robust increase in fluorescence in axons transfected with Syt1-SEP (**Fig. 3e,f**) or VAMP2-SEP (**Supplementary Fig. 2i**), and in axons of untransfected neurons without detectable SEP. We found that the decay kinetics after stimulation was faster for the VO-antibody conjugate (8.8 ± 0.5 s) than the Syt1-SEP (12.2 ± 0.9 s, n = 41; paired t-test p < 0.0001, **Fig. 3f**) and VAMP2-SEP (**Supplementary Fig. 2j**), suggesting a difference in overexpressed vs endogenous protein behavior after SV exocytosis^18^. Remarkably, the fluorescence transients were substantially higher in untransfected than in transfected neurons (**Fig. 3f**, **Supplementary Fig. 2i**), perhaps stemming from steric hindrance of the overexpressed SEP proteins or through quenching of the two fluorophores.

In summary, we have developed new ‘semisynthetic’ pH-sensitive proteins that allow for the imaging of synaptic vesicle fusion events in living cells. This sensor system combines the highly tunable properties of small molecule fluorophores with the specificity of self-labeling tags or antibodies. The SNAP-tag-based system constitutes the first genetically encoded long-wavelength pH sensor with similar or better performance than SEP in different cell types. The antibody-based pH sensor allows for imaging of vesicle fusion events without the need for overexpression of sensor proteins. Addition of other self-labeling or epitope tags by genome editing methods could allow cell- and protein-specific labeling without the need for overexpression^19^. Future improvements of both the protein and the dye within this semi-synthetic scaffold should further enable imaging of this key biological process in increasingly complex systems.

## METHODS

Methods, including statements of data availability and any associated accession codes and references, are available in the online version of the paper.

## ACKNOWLEDGMENTS

We thank the cell culture core facility of IINS for preparing neuronal cultures, and Marie-Paule Strub (NIH) for assistance with molecular biology. This work was supported by the Agence Nationale de la Recherche (to D.P.), the ERC (to D.C.), the Intramural Research Program of the National Heart, Lung, and Blood Institute, NIH (to J.W.T.), and the Howard Hughes Medical Institute (to L.D.L.). M.M. is the recipient of a Marie Sklodowska-Curie Individual Fellowship (IF) under the Horizon 2020 Program (H2020) of the European Commission.

## AUTHOR CONTRIBUTIONS

M.M. performed and analyzed experiments on neurons. A.S. performed experiments on PC12 cells, and A.S. and J.W.T. analyzed data. J.B.G. performed organic synthesis. T.D.G. prepared and analyzed protein conjugates. L.D.L. performed spectroscopy. M.M., L.D.L., J.W.T. and D.P. wrote and all the other authors edited the manuscript.

## ONLINE METHODS

### Organic synthesis

Experimental procedures and characterization data for all new compounds can be found in the Supplementary Note.

### UV-Vis and fluorescence spectroscopy

Spectroscopy was performed using 1-cm path length, 3.5-mL quartz cuvettes or 100-μL quartz microcuvettes Starna Cells. All measurements were taken at ambient temperature (22 ± 2 °C). Absorption spectra were recorded on a Cary Model 100 spectrometer (Varian), and fluorescence spectra were recorded on a Cary Eclipse fluorometer (Varian). The pKa values for compounds **1**, **2** and **3** were determined in buffers containing 150 mM NaCl and 10 mM buffer. The following buffer systems were used: citrate (pH 4.0–6.2); phosphate (pH 5.8– 8.0); tris (pH 7.8–9.0); carbonate (pH 9.2–10.0). Fluorescence values were read on 500 nM samples (n = 3) and fitted to a sigmoidal dose response curve (variable slope) using GraphPad Prism software. Samples for visual inspection containined 5 pM **1** or **2** in citrate (pH 5.6) or phosphate (pH 7.4) buffer.

### Protein chemistry

SNAP-tag protein (NEB) was labeled with excess CFl SNAP-tag ligand (**4**) in PBS containing 1 mM DTT. The protein concentrate was purified and concentrated using a Ziba spin desalting column (7K MWCO; ThermoFisher). The protein sample was analyzed by gel electrophoresis using a NuPAGE 4–12% Bis-Tris polyacrylamide gel (ThermoFisher) and imaged using a Typhoon Trio+ scanner (GE Healthcare); the gel was also stained with Coomassie Plus (ThermoFisher) and compared to a Mark 12 protein standard ladder (ThermoFisher). Absorbance measurements were taken in citrate (pH 5.6) or phosphate (pH 7.4) buffers described above.

### Plasmid constructs

VAMP2-SEP and synaptotagmin1-SEP were kind gifts from Jürgen Klingauf. GCaMP6f was obtained from Addgene (#40755). VaChT-SNAP and VaChT-pHuji were created by replacing the open reading frame of SEP for either SNAP-tag (New England Biolabs) or pHuji^5^ in the VAChT-SEP construct^15^. VAMP2-pHuji was created by swapping the SEP gene for pHuji in VAMP2-SEP sensor construct^2^.

### Cell culture and transfections

PC12 cells were grown in DMEM containing 4 mM L-glutamine, supplemented with 5% fetal bovine serum, 5% horse serum, and 1% penicillin/streptomycin, at 37 °C in 5% CO_2_ Cells were plated onto 20-mm round poly-D-lysine coated glass cover-slips and transfected approximately 24–48 h later with 1 μg VAChT-SEP, VAChT-SNAP or VAChT-pHuji plasmids using Lipofectamine 2000 (Invitrogen) following the manufacturer’s protocol. Cells were imaged 24 h after transfection.

Dissociated hippocampal neurons from E18 rat embryos of either sex were prepared as described previously^20^ at a density of 300,000 cells per dish on poly-D-lysine-coated coverslips, and maintained at 37 °C and 5% CO2 in Neurobasal medium supplemented with 2 mM glutamine and 1× NeuroCult SM1 Neuronal supplement (STEMCELL Technologies). Transfection of VAMP2-SEP, VAMP2-pHuji, VAMP2-SNAP, GCaMP6f and Syt1-SEP was performed at 6 days in vitro (DIV) by a modified calcium phosphate transfection procedure^21^. Experiments were carried out at 20–25 DIV.

### Antibody labeling

Mouse monoclonal antibodies against the luminal part of Synaptotagmin 1 (100 μg, Synaptic Systems 105 311) were coupled in carbonate buffer (pH 9) to 3.4 eq VOAc_2_-NHS ester (**6**; solubilized in DMSO) for 1 h. VO was then deacetylated by hydroxylamine (150 mM) overnight at 4 °C. Labelled antibodies were purified on a size exclusion column and eluted in PBS at a final concentration of 0.1 μΜ. The degree of labelling was estimated to be 6.8 dyes per antibody by absorption spectroscopy.

### Cell staining with CFl and VO ligands

PC12 cells were washed with Fluorobrite media (Fluorobrite DMEM (Invitrogen) supplemented with 5% fetal bovine serum, 5% horse serum, 4 mM L-glutamine, 1% pyruvate, and 1% penicillin/streptomycin), and incubated with 6 μM CFl-SNAP-tag ligand (**4**) or VO-SNAP-tag ligand (**5**) in Fluorobrite media for 3 h at 37 °C. Cells were subsequently washed thoroughly, and incubated in media for an additional 2 h before imaging. SVs were labelled with CFl-SNAP-tag ligand (**4**), VO-SNAP-tag ligand (**5**) and anti-Syt1-VO antibody. Neurons were exposed to 10 μM CFl-SNAP-tag ligand (**4**) or VO-SNAP-tag ligand (**5**) for 1 h in conditioned culture medium, washed thoroughly and then placed back in conditioned culture medium for at least 30 min before imaging. Neurons were incubated with 10 nM anti-Syt1-VO in a 37°C incubator for 3 h in a carbonate buffer containing 105 mM NaCl, 20 mM KCl, 2.5 mM CaCl_2_, 1 mM MgCl_2_, 10 mM glucose, 18 mM NaHCO_3_. Cells were then washed three times before imaging.

### Live cell fluorescence imaging and analysis of PC12 cells

Cells were imaged in buffer containing 130 mM NaCl, 2.8 mM KCl, 5 mM CaCl_2_, 1 mM MgCl_2_, 10 mM HEPES and 10 mM glucose. pH was adjusted to 7.4 with 1 N NaOH. The stimulation buffer contained 50 mM NaCl, 105 mM KCl, 5 mM CaCl_2_, 1 mM MgCl2, 10 mM HEPES and 1 mM NaH2PO_4_. pH was adjusted to 7.4 with 5 M KOH. Experiments were carried out at 25 °C. TIRF microscopy was done as previously described^14,22^. Cells were imaged on an inverted fluorescent microscope (IX-81, Olympus), equipped with a X100, 1.45 NA objective (Olympus). 488 nm and 561 nm lasers (Melles Griot) were combined and passed through a LF405/488/561/635 dichroic mirror. The laser was controlled with an acousto-optic tunable filter (Andor). Emitted light was separated using a 565 DCXR dichroic mirror on the image splitter (Photometrics), and projected through 525Q/50 and 605Q/55 filters onto the chip of a EM-CCD camera. Image acquisition was done using the Andor IQ2 software. Images were acquired sequentially with alternate 488nm and 561nm excitation at 100 ms exposure. The red and green images were aligned post-acquisition using projective image transformation as described before^14,22^. Before experiments, 100 nm yellow-green fluorescent beads (Invitrogen) were imaged in the green and red channels, and superimposed by mapping bead positions.

Image analysis was performed using Metamorph (Molecular Devices) and custom scripts on MATLAB (Mathworks). The co-ordinates of the brightest pixel in the first frame of each fusion event in the green channel was identified by eye, and time was normalized to 0s. A circular ROI of 6 pixels (∼ 990 nm) diameter and a square of 21 pixels (∼ 3.5 μm) were drawn around the fusion co-ordinates. The average minimum pixel intensity in the surrounding square from 5 frames before fusion was subtracted from the intensity in the circular ROI, and the values were normalized to the frame before fusion in the green and red channels.

### Live cell fluorescence imaging and analysis of hippocampal neurons

The experiments with neurons were carried out in a buffer solution containing 120 mM NaCl, 5 mM KCl, 2 mM CaCl_2_, 2 mM MgCl_2_, 5 mM glucose, 10 mM HEPES adjusted to pH 7.4 and 270 mOsmol/l. Neurons were stimulated by electric field stimulation (platinum electrodes, 10 mm spacing, 1 ms pulses of 50 mA and alternating polarity at 20 Hz) applied by constant current stimulus isolator (SIU-102, Warner Instruments) in the presence of 10 μΜ 6-cyano-7-nitroquinoxaline-2,3-dione (CNQX) and 50 μΜ D,L-2-amino-5-phosphonovaleric acid (AP5) to prevent recurrent activity.

Experiments were performed on an inverted microscope (IX83, Olympus) equipped with an Apochromat N oil 100^×^ objective (NA 1.49). Images were acquired with an electron multiplying charge coupled device camera (QuantEM:512SC; Roper Scientific) controlled by MetaVue7.1 (Roper Scientific). Samples were illuminated by a 473-nm laser (Cobolt) for green imaging, as well as by a coaligned 561-nm laser (Cobolt) for red imaging. Emitted fluorescence was detected after passing filters (Chroma Technology Corp.): 595/50 nm for pHuji, CFl and VO imaging, and 525/50 nm for SEP/GFP imaging. Simultaneous dual color imaging was achieved using a DualView beam splitter (Roper Scientific). To correct for x/y distortions between the two channels, images of fluorescently labeled beads (Tetraspeck, 0.2 μm; Invitrogen) were taken before each experiment and used to align the two channels^5^. Time lapse images were acquired at 1 or 2 Hz with integration times from 50 to 150 ms.

Image analysis was performed with custom macros in Igor Pro (Wavemetrics) using an automated detection algorithm as described previously^21^. The image from the time series showing maximum response during stimulation was subjected to an “à trous” wavelet transformation. All identified masks and calculated time courses were visually inspected for correspondence to individual functional boutons. The intensity values were normalized to the ten frames before stimulation in the green and red channels. Photobleaching in the red channels was corrected using an exponential decay fit applied on the non-responsive boutons. All data are represented as mean ± s.e.m. of n experiments.

---

**Supplementary Figure 1, related to Figure 2:**
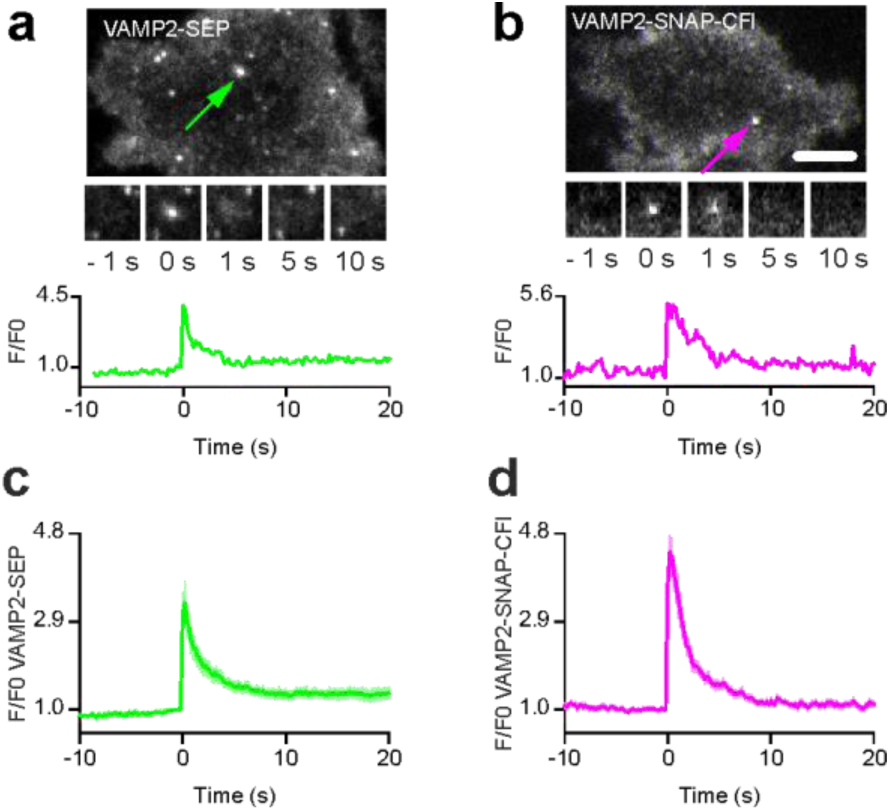
Detection of exocytosis events in PC12 cells with VAMP2 fusion proteins. (**a**) Fluorescence image of a PC12 cell expressing VAMP2-SEP with the inset indicated with the green arrow (top). Time-lapse and fluorescence of the exocytosis event (bottom). (**b**) Same as (a) for a PC12 cell expressing VAMP2-SNAP incubated with VO-SNAP-tag ligand. (**c**,**d**) Average relative fluorescence of events recorded in cells expressing VAMP2-SEP (d, 34 events in 3 cells) or VAMP2-SNAP-CFl (e, 37 events in 5 cells).

**Supplementary Figure 2, related to Figure 3:**
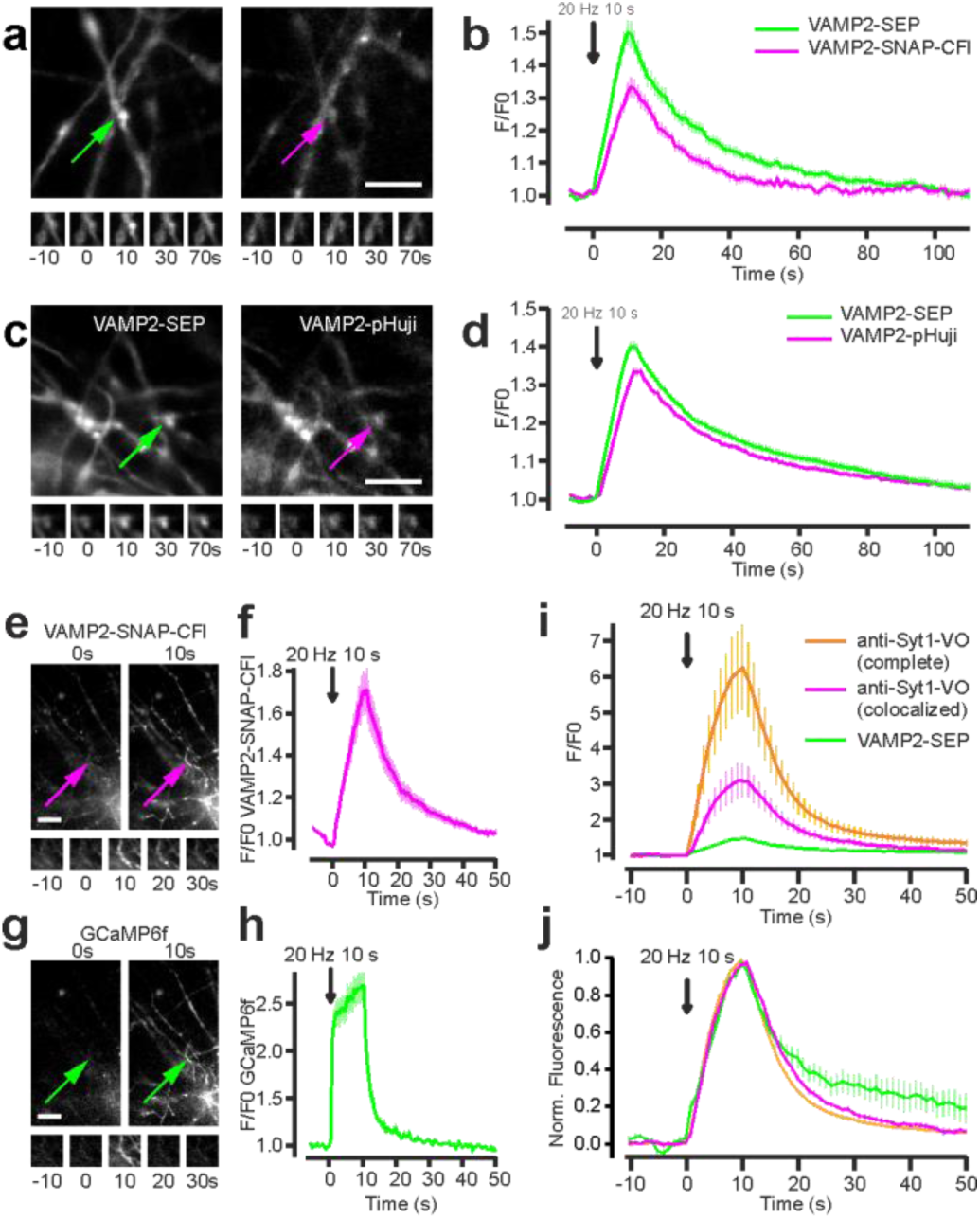
Detection of synaptic vesicle exocytosis and recycling in hippocampal neurons. (**a**) Fluorescence images of hippocampal axons expressing VAMP2-SEP (left) and VAMP2-SNAP-CFl (right) with the insets indicated with arrows (top). Time-lapse of the exocytosis events (bottom). Scale bar: 5 μm. (**b**) Average VAMP2-SEP (green) and VAMP2-SNAP-CFl (magenta) fluorescence signals in response to field stimulation for 10 s at 20 Hz. Fluorescence decay constant after stimulation was 17.8 ± 1.5 s for SEP and 15.6 ± 1.0 s for SNAP-CFl (n = 25). (**c**) Fluorescence images of hippocampal axons expressing VAMP2-SEP (left) and VAMP2-pHuji (right) with the insets indicated with arrows (top). Time-lapse of the exocytosis events (bottom). (**d**) Average VAMP2-SEP (green) and VAMP2-pHuji (magenta) fluorescence signals in response to field stimulation for 10 s at 20 Hz. Fluorescence decay constant after stimulation was 16.2 ± 0.8 s for SEP and 19.6 ± 0.9 s for pHuji (n = 60). Scale bar: 5 μm. (**e-h**) Simultaneous detection of synaptic vesicle cycle and calcium transients in hippocampal neurons. (**e,g**) Fluorescence images of hippocampal axons expressing VAMP2-SNAP-CFl (**e**) and GCaMP6f (**g**). Scale bar: 10 μm. (**f,h**) Average VAMP2-SNAP-CFl (**f**) and GCaMP6f (**h**) fluorescence signals in the same boutons in response to field stimulation for 10 s at 20 Hz. n = 26. (**i**) Detection of synaptic vesicle cycle using endogenous labels. Average anti-Syt1-VO (magenta) and VAMP2-SEP (green) fluorescence signals colocalized in the same boutons, and average anti-Syt1-VO fluorescence signals present in the complete neuronal population (orange) in response to field stimulation for 10 s at 20 Hz. (**j**) Average normalized traces corresponding to the recordings in (n). Fluorescence decay after stimulation was 10.6 ± 1.9 s for VAMP2-SEP and 7.9 ± 0.4 s for anti-Syt1-VO co-localized with VAMP2-SEP, and 6.5 ± 0.2 for all boutons (n = 10 fields). Data are represented as mean ± s.e.m.

